# Cryptic prophages within a *Streptococcus pyogenes* genotype *emm*4 lineage

**DOI:** 10.1101/2020.05.19.103838

**Authors:** Alex Remmington, Samuel Haywood, Julia Edgar, Claire E. Turner

**Affiliations:** Department of Molecular Biology & Biotechnology, The Florey Institute, University of Sheffield, Sheffield, UK; Department of Molecular Biology, Princeton University, USA

**Keywords:** group A *Streptococcus*, bacteriophage, superantigen, DNase, SpyCI, DNA mismatch repair

## Abstract

The major human pathogen *Streptococcus pyogenes* shares an intimate evolutionary history with mobile genetic elements, which in many cases, carry genes encoding bacterial virulence factors. During recent whole genome sequencing of a longitudinal sample of *S. pyogenes* isolates in the United Kingdom, we identified a lineage within *emm*4 that clustered with the reference genome MEW427. Like MEW427, this lineage was characterised by substantial gene loss within all three prophage regions, compared to MGAS10750 and isolates outside of the MEW427-like lineage. Gene loss primarily affected lysogeny, replicatory and regulatory modules, and to a lesser and more variable extent, structural genes. Importantly, prophage-encoded superantigen and DNase genes were retained in all isolates. In isolates where the prophage elements were complete, like MGAS10750, they could be induced experimentally, but not in MEW427-like isolates with degraded prophages. We also found gene loss within the chromosomal island SpyCIM4 of MEW427-like isolates, although surprisingly, the SpyCIM4 element could not be experimentally induced in either MGAS10750-like or MEW427-like isolates. This did not, however, appear to abolish expression of the mismatch repair operon, within which this element resides. The inclusion of further *emm*4 genomes in our analyses ratified our observations and revealed an international *emm*4 lineage characterised by prophage degradation. Intriguingly, the USA population of *emm*4 *S. pyogenes* appeared to constitute predominantly MEW427-like isolates, whereas the UK comprised both MEW427-like and MGAS10750-like strains. The degradation and cryptic nature of these elements may have important phenotypic ramifications for *emm*4 *S. pyogenes* and the geographical distribution of this lineage raises interesting questions on the population dynamics of the genotype.

**Data summary:** All raw sequence data used in this study has been previously published and was obtained from NCBI short read archive. Accession numbers and citations for the genome data for each individual isolate is provided in Supplementary Table 1.

## Introduction

*Streptococcus pyogenes* (also known as the Lancefield Group A *Streptococcus* or GAS) is a globally distributed human pathogen (Efstratiou and Lamagni, 2016). The spectrum of infections caused by *S. pyogenes* is broad, with manifestations ranging from the relatively mild and self-limiting tonsillitis and scarlet fever, to the more severe, invasive and potentially life threatening, notably necrotising fasciitis and streptococcal toxic shock syndrome (Carapetis *et al*., 2005). *S. pyogenes* isolates are classified into *emm* genotypes in accordance with the 5’ hypervariable region of the *emm* gene, which encodes the surface protein M (Facklam *et al*., 1999; Mcmillan *et al*., 2013).

Most *S. pyogenes* genotypes are lysogenized by at least one prophage, though it is not at all uncommon for certain genotypes to carry several such elements integrated into the bacterial chromosome, or indeed none (McShan and Nguyen, 2019). As is the case for a number of other bacterial pathogens, the lysogenic bacteriophages in this species often are associated with bacterial virulence factors. Eight of the thirteen superantigens in *S. pyogenes (speA, speC, speH, speI, speK, speL, speM* and allelic variants thereof), are prophage-associated (Proft *et al*., 2003). In addition, prophage in this species often carry genes encoding a potential six secreted nucleases (*sdn, spd1, spd3, spd4, sda, sdaD2*, and allelic variants thereof), and a secreted A_2_ phospholipase, denoted *sla* (Beres *et al*., 2004; Remmington and Turner, 2018). The *S. pyogenes* chromosomal islands (SpyCI) are similar but distinct mobile genetic elements which are more closely related to the *Staphylococcus aureus* pathogenicity islands (SaPIs) (Novick *et al*., 2010; Nguyen and McShan, 2014). In the *emm*1 isolate SF370, SpyCI-M1 is integrated within the DNA mismatch repair (MMR) operon, separating *mutS* from *mutL* and other downstream repair associated genes, preventing co-transcription of these genes from a promoter upstream of *mutS* (Scott *et al*., 2008). A similar element was also identified in the genomes of isolates belonging to other genotypes, including the *emm*4 MGAS10750 (Scott *et al*., 2012). With the exception of *emm*5 isolate Manfredo, the presence of a SpyCI element within the MMR operon was associated with increased mutation rates, indicating the disruption of the MMR-associated gene transcription (Scott *et al*., 2008; Scott *et al*., 2012). Dynamic integration and excision of SpyCIM1 in SF370 was detected in response to bacterial growth; excising during earlier growth phases to permit transcription of the MMR operon, but remaining integrated at later stages, abolishing transcription and promoting a transient, more mutable phenotype (Scott *et al*., 2008; Scott *et al*., 2012). Intriguingly, curing of SpyCIM1 had a dramatic effect on global transcription, including the expression of a number of well characterised virulence factors (Hendrickson *et al*., 2015). Together, these studies effectively describe a mechanism of genetic regulation in *S. pyogenes* that is dependent on integration and excision of phage-like elements.

In recent years, whole genome sequencing (WGS) has provided valuable insight into the population biology of this pathogen, illuminating population dynamics and bringing clarity to outbreak investigations and epidemiological shifts. Often, changes in infection character, population structure and disease incidence have been associated with prophage being acquired and/or lost from the population, and with them, cognate bacterial virulence factors, particularly the streptococcal superantigens and DNases (Banks *et al*., 2002; Beres *et al*., 2004; Tse *et al*., 2012; Ben Zakour *et al*., 2015; Turner *et al*., 2015; Walker, 2015; Al-Shahib *et al*., 2016; Afshar *et al*., 2017). Genotype *emm*4 *S. pyogenes* constitute a major *emm*-type in high income countries globally (Efstratiou and Lamagni, 2016) that is capable of causing both superficial and invasive disease (Jaggi *et al*., 2005; Luca-Harari *et al*., 2009; Whitehead *et al*., 2011; Flores *et al*., 2019). This genotype is genetically acapsular (Flores *et al*., 2012), frequently associated with outbreaks of scarlet fever (Yan *et al*., 2003; Silva-Costa *et al*., 2014; Turner *et al*., 2016; Chalker *et al*., 2017) and occasionally has superseded in incidence the consistently dominant *emm*1 genotype, in some parts of the world (Whitehead *et al*., 2011; Kim *et al*., 2019). Typically, strains belonging to this genotype are host to three prophage elements; Φ10750.1, Φ10750.2 and Φ10750.3, encoding the streptococcal superantigen *speC* and the DNase *spd1*, the DNase *spd3*, and another streptococcal superantigen *ssa*, respectively (Beres *et al*., 2006; Henningham *et al*., 2014; Galloway-Peña *et al*., 2016; Jacob *et al*., 2016; Turner *et al*., 2019). It was recently reported that in an *emm*4 population from Houston, Texas USA, the majority of isolates carried a novel chimeric *emm* gene, formed by the fusion of the 5’ end of *emm*4 with the 3’ end of the downstream gene *enn*. This chimeric *emm*4 gene was also found in the closely related reference *emm*4 genome MEW427 but not the more distantly related MGAS10750 (DebRoy *et al*., 2018).

Here, we reveal that within an *emm*4 population from England is a lineage that is characterised by marked gene loss within prophage encoding regions, although associated superantigen and DNase genes remain intact. This lineage was also identified in a wider international population. The gene loss has rendered the prophages immobile, unable to replicate extra-chromosomally and are therefore cryptic prophages.

## Results

### Gene loss within each of three prophage regions

We recently undertook WGS of 344 invasive *S. pyogenes* isolates, collected by the British Society for Antimicrobial Chemotherapy (BSAC) from across England during 2001-2011, and identified ten *emm*4 isolates within this collection (Turner *et al*., 2019). Within this small population of *emm*4 was a lineage of five isolates that were genetically related to the reference strain MEW427 (Jacob *et al*., 2016). Four of the remaining five isolates were more closely related to the reference strain MGAS10750 (Beres *et al*., 2007) (Figure 1). Gubbins analysis identified regions of predicted recombination, common to the lineage containing MEW427 and five BSAC MEW427-like isolates, but absent in MGAS10750 and the other five BSAC isolates. Four of the regions of predicted recombination were within each of the three prophages (Φ10750.1, Φ10750.2, Φ10750.3) and the chromosomal island SpyCI (Φ10750.4). Comparison of the three prophage regions between the two completed reference strains MGAS10750 and MEW427 actually identified a varying level of gene loss within these regions in the MEW427 genome compared to the MGAS10750 genome (Figure 2). To determine if an equivalent gene loss had occurred in the genomes of the BSAC isolates, the presence or absence of each gene within the MGAS10750 prophages was confirmed by BLAST analysis of the *de novo* assembled BSAC genome sequence data (Figure 2). In all isolates, the superantigen and DNase genes associated with each of the prophages were present and shared 100% DNA identity between all ten BSAC isolates and the reference strains.

**Figure 1.**
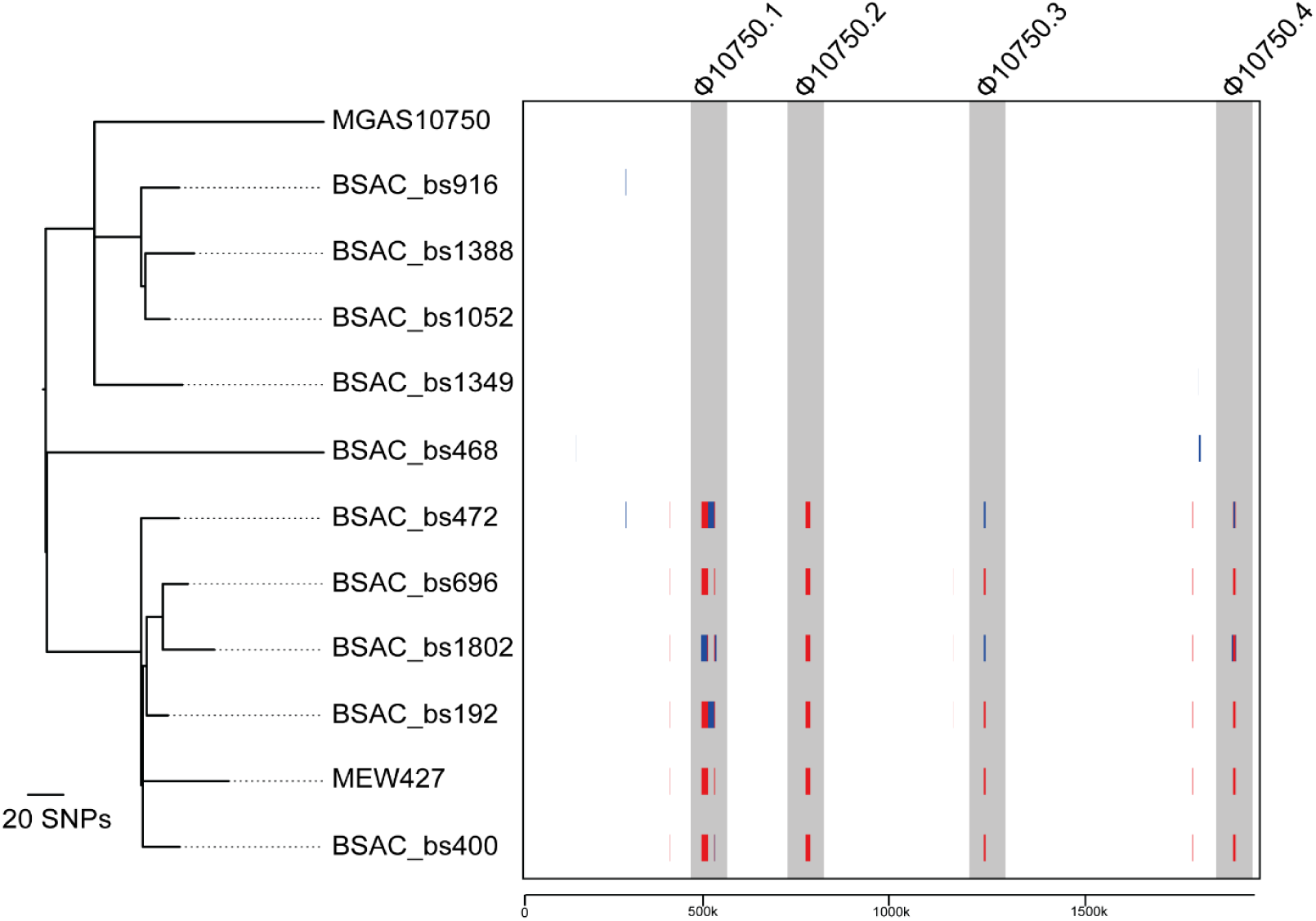
Recombination in prophage regions of isolates clustering with MEW427. Sequence data for all ten BSAC isolates were mapped to the reference strain MGAS10750 along with a second reference strain MEW427. Gubbins analysis on the alignment identified predicted regions of recombination in five BSAC isolates and MEW427 that form a separate lineage to the other five BSAC isolates and MGAS10750. Four of these regions were within the three prophages (Φ10705.1, Φ10705.2, Φ10705.3) and the SpyCI element (Φ10705.4) (shaded grey). Phylogenetic tree was generated from 556 polymorphic sites following removal of regions of predicted recombination. Scale bar; number of SNPs. Scale below boxed region represents genomic position relative to MGAS10750.

**Figure 2.**
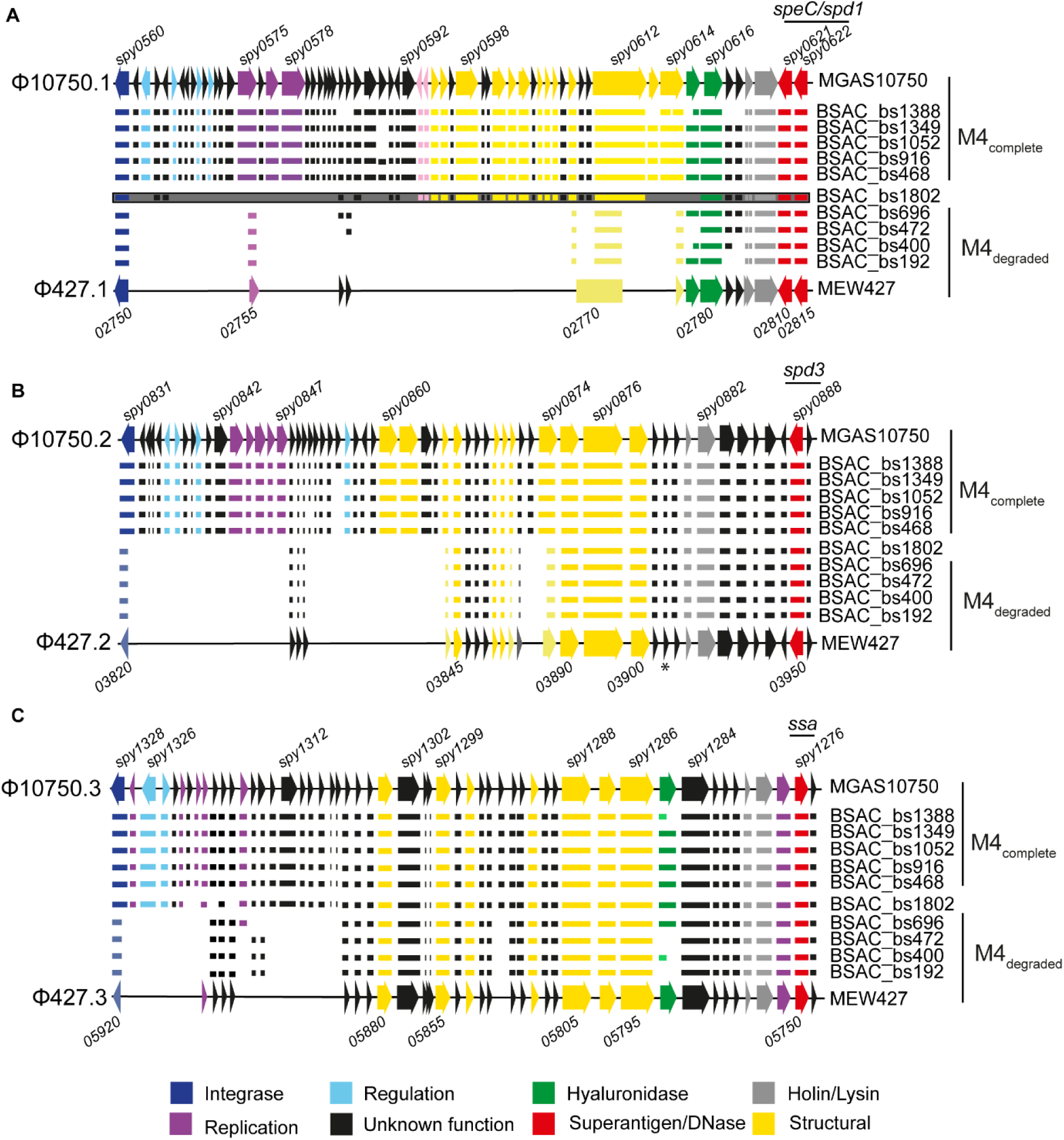
Varying levels of gene loss within prophage regions. The majority of genes (represented by arrows) present in the three prophages (**A**) Φ10705.1, (B) Φ10705.2 and (**C**) Φ10705.3 of MGAS10750 were also present in the five BSAC strains BSAC_bs468, BSAC_bs916, BSAC_bs1052, BSAC_bs1379 and BSAC_bs1388 (M4_complete_). However, fewer genes were present in the corresponding prophage regions (**A**) Φ427.1, (B) Φ427.2 and (**C**) Φ427.3 in MEW427 and the MEW427-like BSAC isolates BSAC_bs192, BSAC_bs400, BSAC_bs472 and BSAC_bs696 (M4_degraded_). The genome of BSAC_bs1802 had more prophage-associated genes present than the other M4_degraded_ isolates, and the prophage associated with *speC*/*spd1* appeared to be different to Φ10705.1/ Φ427.1 (represented by a grey box), with some homologous genes but assembled with other genes that were not found in MGAS10750 or MEW427. Gene presence in each strain is represented by corresponding horizontal lines. Some genes in MEW427 and M4_degraded_ isolates were truncated (shown in lighter colours), for example the integrase genes of Φ427.2 and Φ427.3 (lighter shade of blue and shorter arrow/line). Within Φ427.1, the annotated gene *02770* actually comprised sections of *spy609, spy0612* and *spy614* of MGAS10750. In all isolate genomes and in MEW427, *spy0619* of Φ10705.1 was divided into two genes. Locus numbers for some genes are provided above and below; MEW427 is annotated in increments of 5. *; an additional gene that was present but not included in the reference annotation. Colours represent predicted gene function according to the key.

In MGAS10750, Φ10750.1, associated with *speC/spd1*, comprised 63 genes, while in reference genome MEW427, this element (Φ427.1) comprised just 14, equivalent to ∼70% gene loss. In MEW427-like lineage associated *emm*4 isolates in our collection, with the exception of BSAC_bs1802, we found similar patterns of gene presence/absence (Figure 2A). All five of the other BSAC isolates had all or nearly all (1-6 genes missing) genes that comprise Φ10750.1, indicating this prophage is complete in these isolates (Figure 2A).

The exception was BSAC_bs1802 that, although clustered by core genome SNP analysis within the MEW427-like lineage (Figure 1), showed a different level of Φ10750.1 gene presence/absence. We could not obtain fully assembled sequence data across the prophage region with our short-read sequence data and this region was split over multiple contigs. From our analysis, however, it appeared that BSAC_bs1802 actually carried *speC/spd1*on a different prophage to the rest of the *emm*4 population; this shared a high level of homology to Φ10750.1 at the 3’ integration site near the *speC*/*spd1* genes, but differed at the 5’ end downstream of the integrase gene, with the presence of other non-Φ10750.1 genes. This was confirmed with two methods of *de novo* assembly. The integration site remained the same as in MGAS10750/MEW427.

We did not detect any non-Φ10750.1 prophage genes within this region in the other four MEW427-like isolate genomes, as we did with BSAC_bs1802, but we did identify genes found in Φ427.1 that were altered by the gene loss; MEW427-*02755* which is the truncated equivalent of MGAS10750-*spy0575*, and MEW247-*2770* which is a fusion of regions of MGAS10750-*spy609, spy612*, and *spy614* (Figure 2A). This indicated that within these four BSAC isolate genomes, there was the same (or very similar) degraded prophage as Φ427.1 in MEW427. Genes predicted to be involved in regulation and replication of the prophage element were absent in Φ427.1 and in the four MEW427-like BSAC isolates. All predicted structural genes were also absent from the genomes of these isolates, suggesting this prophage would be unable to form bacteriophage particles.

The *spd*3-associated prophage Φ10750.2, comprised 59 genes in MGAS10750 and a similar pattern was observed in five BSAC isolates (Figure 2B). However, this element in MEW427 and corresponding elements in all five MEW427-like BSAC isolates comprised only 29 genes, a loss of ∼50%. We obtained complete *de novo* assembly across this prophage region in all five MEW427-like isolates confirming the same pattern of gene loss. This included loss of all regulation and replication elements and some structural genes. We found that the first 184/1140 bases of the integrase gene were missing, likely to render the gene non-functional, predicting that this prophage would be unable to excise from the chromosome or integrate.

The *ssa* associated prophage Φ427.3, demonstrated the least gene loss compared to Φ10750.3 (∼26%), maintaining the majority of structural genes, although genes predicted to be involved in replication and regulation were absent in MEW427 and MEW427-like BSAC isolates, except for BSAC_bs1802 (Figure 2C). As with Φ10750.1, we could not determine the complete sequence for the *ssa-*associated phage from our sequence data but we could determine the integration sites to be the same as for Φ10750.3 and we did not identify the presence of any other non-Φ10750.3 genes. The presence of 87% of Φ10750.3 genes in the genome of BSAC_bs1802 suggested it carried the same *ssa-*encoding prophage as MGAS10750, and that this prophage is near complete. The other four MEW427-like BSAC isolates had a very similar pattern of gene presence/absence to MEW427. The integrase gene was also truncated in MEW427 and the four MEW427-like BSAC isolates, with the loss of the first 548/1143 base pairs. We therefore predict that this prophage would also be unable to excise from the chromosome or to reintegrate.

Overall, five BSAC isolates had 87-100% of the genes of all three prophage genes and were therefore termed M4_complete_ type, like MGAS10750 (as indicated in Figure 2). Four BSAC isolates had fewer than 30%, 50% and 70% of the genes present in Φ10750.1, Φ10750.2, Φ10750.3, respectively, like MEW427, and were therefore termed M4_degraded_. BSAC_bs1802 had a different, potentially hybrid phage associated with *speC/spd1*, and 46% and 87% of Φ10750.2 and Φ10750.3 genes, respectively, and was therefore regarded as only partially degraded.

### Prophage excision potential is abolished in isolates with degraded prophage

With the substantive loss of genes within Φ427.1 and the predicted non-functional integrase genes of Φ427.2 and Φ427.3, we hypothesised that the prophages of M4_degraded_ type BSAC isolates would be unable to excise from the chromosome. Using PCR to detect the integrated prophage genome as well as excised and circularised prophage, we found that all three prophage elements spontaneously excise from the chromosomes of M4_complete_ BSAC isolates, and this excision is enhanced with the addition of mitomycin C (Figure 3). In contrast we were unable to detect excision of the three degraded prophages from the chromosomes of M4_degraded_ type BSAC isolates, even upon addition of mitomycin C. For the partially degraded BSAC_bs1802, clear excision of the *speC/spd1*-associated prophage was detected as was *ssa-* associated Φ10750.3/427.3 but not the *spd3*-associated Φ10750.2/427.2. This was expected as the integrase of Φ10750.2/427.2 was truncated in BSAC_bs1802, like the other MEW427-like isolates.

**Figure 3.**
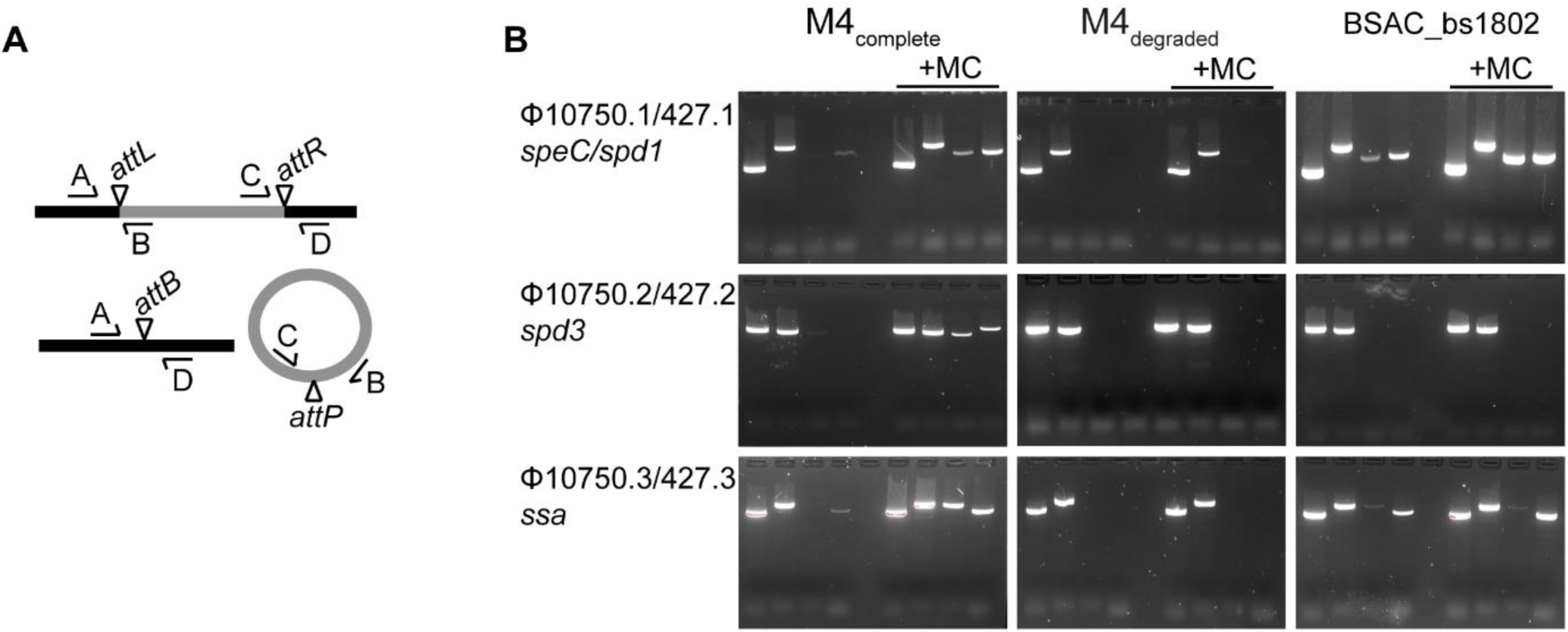
Phage induction in MGAS10750-like isolates but not MEW427-like isolates. **A**. Primers were designed to detected integrated and excised prophage. Primer pairs A+B and C+D spanned the attachment sites *attL* and *attR* formed when the prophage (grey) is integrated into the chromosome (black). Primer pairs A+D and C+B spanned the attachment site *attB* on the bacterial chromosome and *attP* on the prophage, respectively, detecting prophage excision. **B**. All three prophages were detected to be excised from the chromosome of M4_complete_ isolates, which was enhanced by additional of mitomycin C (+MC), as indicated by bands present in all four lanes. No excision was detected for all three prophages in any of the M4_degraded_ isolates, as indicated by bands only in the first two lanes but not the third or fourth lane, even with mitomycin C. The exception was BSAC_bs1802, where excision of Φ10750.1/427.1 and Φ10750.3/427.3 was detected but Φ10750.2/427.2 was not. Representative gels are shown for single isolates out of five M4_complete_ BSAC isolates tested and four M4_degraded_ tested. Primer pairs were used in the following order; A+B (*attL*), C+D (*attR*), A+D (*attB*), C+B (*attP*).

### Gene loss within the SpyCI element

As well as the three prophage regions within MGAS10750, there is an additional mobile genetic element, similar to that previously described for the *emm*1 strain SF370, termed SpyCI (*<u>S</u>. <u>py</u>ogenes* <u>C</u>hromosomal Island). This SpyCI element is integrated between the DNA mismatch repair genes, *mutS* and *mutL* which, in *emm*1 SF370, prevents transcription of *mutL* and disruption of DNA repair. However, it has been shown that during very early exponential growth, this element excises to allow for transcription of *mutL* (Scott *et al*., 2008). In our five BSAC M4_complete_ isolates, this element is identical to that found in MGAS10750, integrated between *mutS* and *mutL*, however it shares 97% DNA identity but only over 55% length with the SpyCI in SF370. Interestingly, like the prophage elements, MEW427 demonstrates some genes loss within the equivalent SpyCI region compared to MGAS10750 (Figure 4). An identical pattern of gene loss was identified in four of the M4_degraded_ BSAC *emm*4 isolates. The exception was BSAC_bs1802, which in fact appear to carry a different SpyCI element that was ∼99% identical to that found in *emm*77 NCTC13742 (Genbank accession: LS483386.1). The SpyCI integrase region in all *emm*4 isolates was identical. PCR was used to detect excision of SpyCI in M4_complete_ and M4_degraded_ *emm*4 isolates, similar to the three prophage regions. We could not detect excision of this element in an M4_complete_ isolate or an M4_degraded_ isolate, even in the presence of mitomycin C (Supplementary Figure 1A). Previous studies have shown that this element excises during early exponential growth (Scott *et al*., 2008), therefore we tested for excision during exponential growth of one M4_complete_ isolate, but still could not detect excision (Supplementary Figure 1B). This was surprising as, when integrated, the element should interrupt the *mutS* and *mutL* operon, preventing the expression of *mutL*. To confirm lack of expression of *mutL*, we extracted RNA from samples cultured for 3 hours, converted the RNA to cDNA and performed semi-quantitative PCR to detect *mutS* and *mutL* transcription. In all samples we detected transcript of both *mutS* and *mutL* suggesting, despite the consistent integration of SpyCI, transcription of *mutL* could still occur (Supplementary Figure 2).

**Figure 4.**
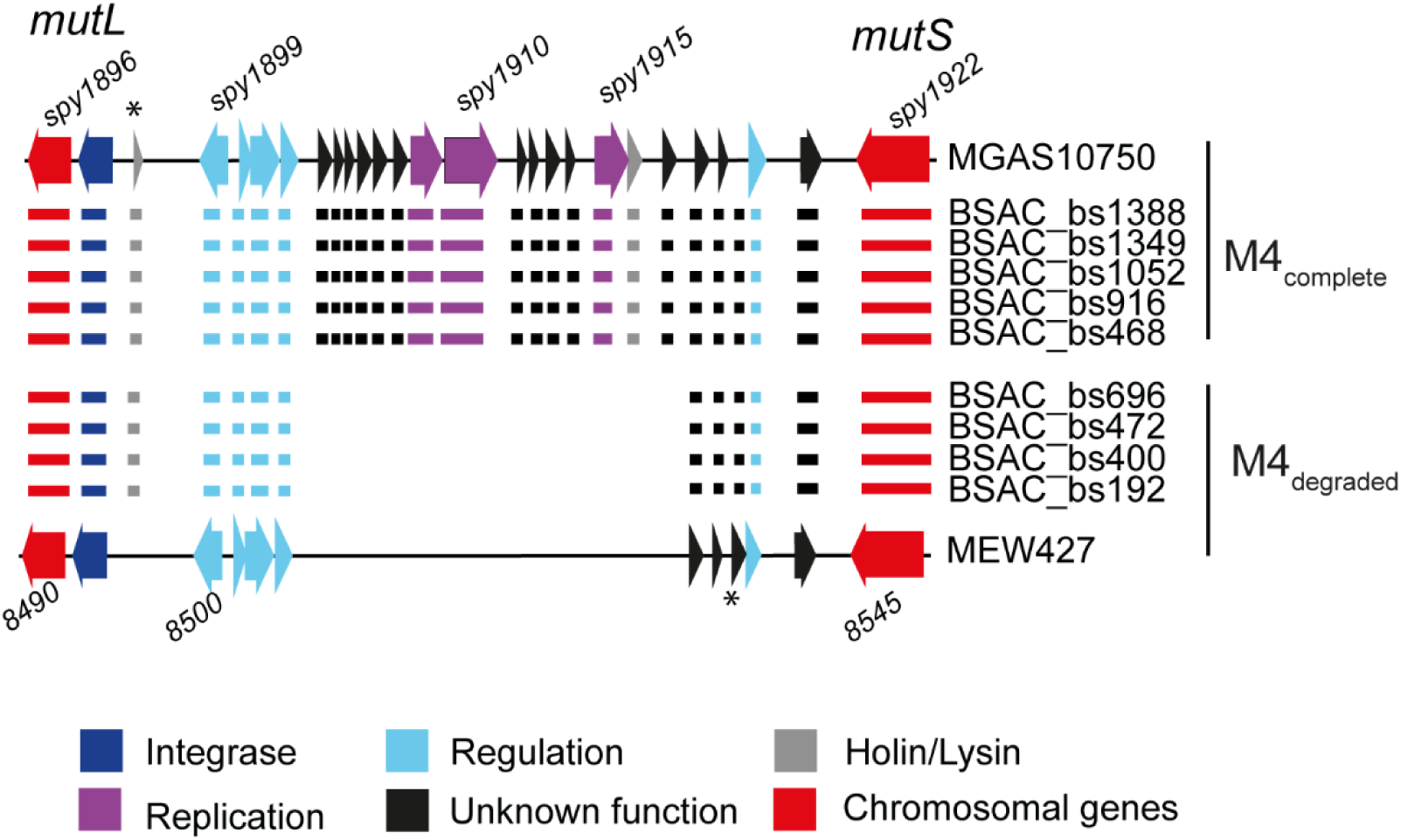
Gene loss within the SpyCI. The SpyCIM4 element is integrated within the MMR operon, between *mutL* and *mutS* (red arrows) and comprised 25 genes at full-length (as in MGAS10750) in the genomes of M4_complete_ isolates. In the genomes of M4_degraded_ isolates, including MEW427, the SpyCI element consists of only 11 genes, representing a ∼56% gene loss. The pattern of degradation was common to all M4_degraded_. The SpyCI of BSAC_bs1802 is not shown, as this isolate contains a different ICE. Locus numbers for some genes are provided above and below; MEW427 is annotated in increments of 5. *; an additional gene that was present but not included in the reference annotation. Colours represent predicted gene function according to the key.

### Prophage-associated gene loss is found in other international *emm*4 isolates

To place our isolates within context of a wider *emm*4 population, we obtained available *emm*4 whole genome sequence data from North America and the UK (n=223) (Supplementary Table 1). Phylogenetic analysis identified two broad lineages associated either with MGAS10750 or with MEW427, as well as a third smaller lineage (Figure 5). BLAST analysis of the genes present in the each of the three MGAS10750 prophages against all *de novo* assemblies (n=223) indicated a varying degree of prophage gene loss for each isolate (Supplementary Figure 3). However, there was an association of fewer prophage genes across all three prophages within a lineage clustering with MEW427 compared to the rest of the population. Although we also identified a varying level of gene loss within our isolates, a consistent feature within MEW427, M4_degraded_ and partially degraded BSAC_bs1802 was a deletion of the 5’ region of the integrase gene in the *spd3* phage Φ10750.2/427.2. All 117 genomes of isolates clustering with MEW427 (shaded region in Figure 5, Supplementary Table 1) had the same deletion within the integrase gene, like MEW427. In contrast, all of the 106 genomes of isolates outside this lineage had a full-length integrase gene, like MGAS10750. We also found within MEW427 and our M4_degraded_ isolates, a deletion of the 5’ end of the *ssa*-associated prophage integrase. This deletion was found in 112/117 genomes of isolates that clustered with MEW427 (shaded region in Figure 5) and complete in the remaining 5 isolates as well as all isolates outside of this lineage. Taken together, it suggests that the *spd3* and *ssa* associated prophages within the majority of isolates clustering within the MEW427-like lineage would not be able to excise from the chromosome and are therefore cryptic.

**Figure 5.**
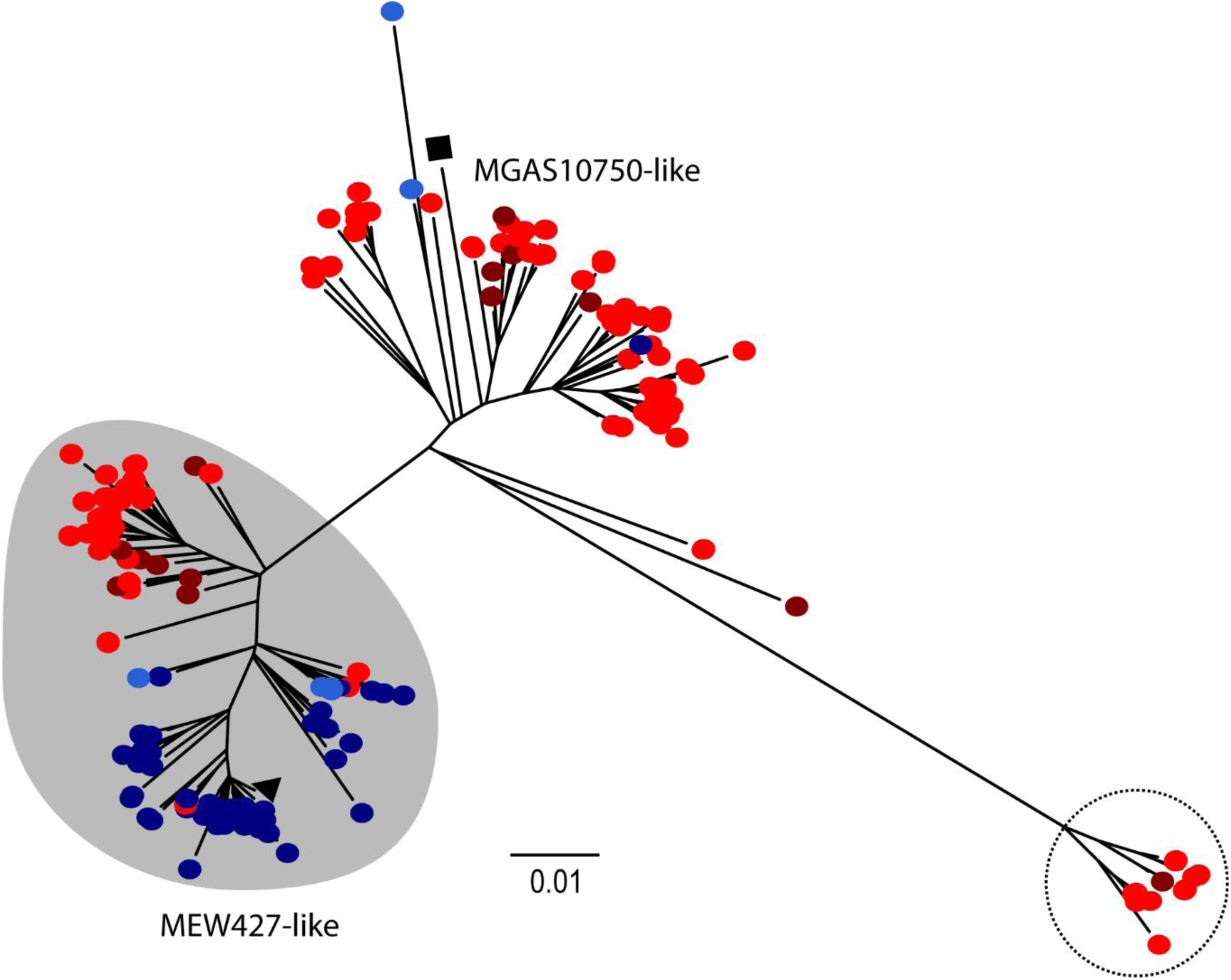
Lineages within the *emm*4 population are associated with prophage gene loss. Genome sequence data from 223 isolates was obtained and mapped against the reference genome MGAS10750 (black square). Additionally, the second reference strain MEW427 was included (black triangle). Data comprised genomes from USA isolates (blue circles, n=48 isolated 2015 (Chochua *et al*. 2017)), Canadian isolates (Light blue circles, n=8 isolated 2013 or unknown date (Athey *et al*. 2014, Athey *et al*. 2016)), UK isolates (red circles, n=153 (Chalker *et al*., 2017; Kapatai *et al*. 2017) isolated 2014/2015), as well as from Cambridgeshire, UK (dark red circles, n=4, 2008-2012, (Turner *et al*., 2017)) and BSAC isolates (dark red circles, n=10, 2001-2008). Isolates in red and dark blue were obtained around a similar time period (2014-2015) but from either the UK (red) or the USA (blue). Two broad lineages exist, either associated with MGAS10750 or with MEW427 (shaded grey). A small lineage was also identified (dotted line circle) and these isolates are all from the UK. Scale bar represents site per substitution.

There was another lineage within the population, comprising 13 UK isolates, for which very few Φ10750.1 genes could be detected, although *speC* and *spd1* genes were present. Prophages Φ10750.2 and Φ10750.3 appeared to be complete with full length integrases. Although we could not fully confirm with the short-read sequence data, it appeared that *speC* and *spd1* were associated with a different prophage in these isolates and not with Φ10750.1 or a degraded form of this prophage. Additionally within the whole population, we also identified two isolates that did not carry the *speC* and *spd1* genes or an associated prophage (Supplementary Table 1).

Interestingly, although UK isolates were divided between the MEW427-like lineage and MGAS10750-like lineage, North American isolates appeared to be mainly restricted to the MEW427-like lineage. Given that the Public Health England isolates (UK) (Chalker *et al*. 2017; Kapatai *et al*. 2017), were collected at a similar time to the USA ABCs collection (2014-2015, (Chochua *et al*., 2017), we compared the numbers of isolates present in each lineage; 57/153 (37%) of PHE isolates were MEW427-like compared to 47/48 (98%) of ABCs isolates.

## Discussion

Genotype *emm*4 *S. pyogenes* is a major *emm*-type causing disease worldwide (Efstratiou and Lamagni, 2016) with the capacity to cause both self-limiting and potentially life-threatening invasive infection (Yan *et al*., 2003; Silva-Costa *et al*., 2014; Turner *et al*., 2016; Chalker *et al*., 2017), and a known association with scarlet fever (Yan *et al*., 2003; Silva-Costa *et al*., 2014; Turner *et al*., 2016; Chalker *et al*., 2017). We have identified a lineage within an international *emm*4 population that is characterised by degradation within integrated prophage genomes. Genetic modules pertaining to replication, regulation and lysogeny were chiefly affected, and to a more variable extent, structural genes necessary for the formation of phage particles. Prophage-associated virulence factors were not affected. We confirmed that, at least in our sample of ten *emm*4 isolates, this degradation resulted in prophage immobility.

The lysogenic prophage of *S. pyogenes* follow a typical lambdoid genomic architecture, with discrete modules dedicated to specific functions (Desiere *et al*., 2001; Canchaya *et al*., 2003; McShan and Nguyen, 2019). The most oblique characteristic of gene loss to prophage-encoding regions in MEW427-like isolates was the replicative and regulatory modules and in two of the three prophages, there were deletions within the integrase genes. From our study, we cannot predict the order in which gene loss occurred, but it seems likely that if a mutation was to occur in the genetic apparatus necessary for excision, selection would presumably act rapidly on those genes remaining that no longer serve a function that is beneficial to the bacterium (Ramisetty and Sudhakari, 2019). As such, the genes or modules remaining raise interesting questions regarding their utility. The retention of prophage-encoded virulence factors is perhaps unsurprising, as multiple lines of scientific investigation have highlighted these genes as conferring a fitness advantage to *S. pyogenes* at the host-pathogen interface (Commons *et al*., 2014; Remmington and Turner, 2018). A potential hypothesis is that genomic inactivation of prophage by the bacterium is part of an ongoing process of domestication, whereby the bacterium may retain genes that are useful, particularly toxin genes, while neutralising the imminent threat of host cell lysis following the induction of the lytic pathway (Ramisetty and Sudhakari, 2019).

Our collection of ten *emm*4 isolates could be divided, within the exception of one isolate, into M4_complete_ and M4_degraded_ as the number of genes present within each of the three prophages reflected that of either the complete (MGAS10750) or degraded (MEW427) reference genome prophages. Complete *de novo* assembly of integrated prophages is notoriously difficult due to the homologous and mosaic nature of these genomes. We were unable to fully assemble these regions in order to identify the exact composition of prophages within each BSAC genome and acknowledge the potential for undetected genetic rearrangements or other prophage elements. This appeared to be the case for BSAC_bs1802 where we identified non-Φ10750.1 genes assembled with the integrase and attachment site of the *speC/spd1* associated prophage, indicative of a hybrid or closely related prophage instead of Φ10750.1. This did not appear to be the case for the four M4_degraded_ isolates, as we detected no genes that did not correspond with Φ10750.1, with the exception of truncated and hybrid genes that were also found in MEW427, which appeared to have arisen through the degradation process, and indicated prophage degradation in these isolates, as opposed to novel prophages. This was also the case for the other two prophages in the M4_degraded_ isolates and BSAC_bs1802. When we extended the analysis to the wider genomic collection there was a high level of variability in the percentage of genes present in each prophage region, but, broadly, gene loss in all three prophages was associated with isolates that clustered with MEW427. All isolates that clustered with MEW427 carried the same deletion leading to a predicted non-functional integrase of the *spd3-*prophage, and the majority (96%) also had the deletion within the *ssa*-prophage integrase. These integrase deletion mutations were not found in isolates outside of this lineage. We predict, therefore, that all isolates belonging to the MEW427-like lineage have at least one immobile and cryptic prophage and it seems likely that all three prophages would be cryptic in many of these isolates.

The phenotypic consequences of prophage degradation are unclear at this stage. An interesting find was the geographical divide whereby all, bar one, isolates from the USA collection (Chochua *et al*., 2017) were found to be MEW427-like, likely representing degraded prophages compared to 37% of UK isolates (Chalker *et al*. 2017; Kapatai *et al*., 2017). This may be related to the over-representation of *emm*4 in the UK associated with scarlet fever (Chalker *et al*. 2017). There is a potential that the prophage, either through direct regulation or excision and replication, may have an impact on the expression of prophage-encoded virulence factors, which include the scarlet fever-associated superantigens. It has previously been shown that induction of toxigenic prophage in *S. pyogenes* and *S. canis* can enhance expression of the cognate virulence factors (Broudy *et al*., 2001; 2002; Banks *et al*., 2003; Ingrey *et al*. 2003). Similar findings have been described in enterohaemorrhagic *E. coli*, wherein toxigenic prophage induction appears to contribute to the pathogenesis of toxin mediated disease (Balasubramanian *et al*., 2019; Zhang *et al*., 2000). It is therefore conceivable that the inactivation of prophage in genotype *emm*4 *S. pyogenes* may represent a progression towards a less virulent phenotype associated with M4_degraded_ isolates, owing to an inability to illicit a gene dosing effect of prophage-encoded virulence factors. It is worth noting, however, that a gene dosing effect does not appear to be necessary in all cases to attain enhanced transcription of these genes (Banks *et al*., 2003).

There are other differences separating the two lineages in addition to the prophage degradation, and these may also impact on the behaviour and success of the lineage. A recent study on *emm*4 isolates from Houston, Texas found isolates with a novel chimeric *emm*, formed from fusion with the downstream *enn* gene (DebRoy *et al*., 2018). This chimeric *emm* was found in MEW427 and MEW427-like isolates but not in MGAS10750. We also identified the chimeric *emm* in our M4_degraded_ BSAC isolates but not in our M4_complete_ BSAC isolates. The chimeric *emm* gene could influence the phenotype of the MEW427-like isolates in addition to the prophage degradation. Chimeric *emm* genes have subsequently been detected in a number of other *emm*-types (Frost *et al*., 2020).

It was interesting to note, that as well as degradation within the three prophage genomes, we also identified degradation within the chromosomal island SpyCI. We expected to find excision of SpyCIM4 from the M4_complete_ lineage isolates during exponential growth or upon exposure to mitomycin C, as observed for SpyCIM1 in SF370 (Scott *et al*., 2008). The excision of SpyCI is essential for the transcription of *mutL*. However, we could not detect the excision of SpyCIM4 from M4_complete_ isolates, nor M4_degraded_ isolates, under any conditions tested. Despite this, transcription of both *mutL* and *mutS* were maintained. It seems that in *emm*4 the MMR operon is not regulated by the integration and excision of the element as has been described in other *emm*-types (Scott *et al*., 2012), although it is possible that low undetectable levels of SpyCI are excised allowing for transcription of *mutL*. We would expect, however, that this would not be the case for the M4_degraded_ isolates, due to the substantial gene loss within this element, yet *mutL* transcription still occurred. The inability to detect SpyCI in its extra-chromosomal conformation in MGAS10750-like isolates, yet apparently retaining a full complement of associated genes relative to the completed reference genome, may indicate that the element can be induced under conditions not explored in the present study, for example in response to antibiotics (Zhang *et al*., 2000; Maiques *et al*., 2006) or factors produced by the host *in-vivo* (Broudy *et al*., 2001, 2002; Banks *et al*., 2002; Broudy and Fischetti, 2003). It is also possible that due to its stably integrated state, the element was subjected to deletion of surplus genes. In a previous study wherein the SpyCI element was cured from an SF370-like *emm*1 isolate, this resulted in dramatic changes to global transcription (Hendrickson *et al*., 2015). Retaining a partial or full-length SpyCI in *emm*4 may therefore serve a function that extends beyond the DNA MMR operon. Additionally, the gene loss within SpyCI, and potentially the other prophage genomes too, could have a wider influence on the phenotype of the bacterium through as yet unidentified transcriptional or even post-transcriptional regulatory mechanisms.

It seems likely that prophage degradation exists in other genotypes as well as *emm*4. Short-read whole genome sequencing technologies are not ideal for the study of prophages as it can be difficult to assemble and to determine gene composition. There has been evidence for prophage-associated genome rearrangements (Nakagawa *et al*., 2003) and hybridisation of different prophages (Davies *et al*. 2015) in *S. pyogenes*. These events may not have been identified using short-read sequence technology. The advancement of long-read sequencing is beginning to allow us to explore the streptococcal genome to a greater extent, with the possibility for more detailed prophage-based research. There is great potential for prophages to drive evolutionary changes within the streptococcal population and their mechanisms of impact could range from simple transfer of virulence factors through to local and genome-wide transcriptional control and even direct phenotypic influences through activation, particle formation and cell lysis. We need to expand our knowledge and research in this area to fully appreciate the role prophages play in the population of streptococci and other bacterial species.

## Materials and Methods

### Bacterial strains

All *S. pyogenes* isolates were grown on Columbia agar supplemented with 5% defibrinated horse blood. For liquid culture, all isolates were grown statically in Todd-Hewitt Broth at 37°C supplemented with 5% CO_2_. *Emm*4 isolates were selected from a collection of 344 *S. pyogenes* bloodstream infection isolates submitted to the British Society for Antimicrobial Chemotherapy from 11 geographical locations within the United Kingdom (Turner *et al*., 2019).

### Genome sequence analysis

*De novo* assembly for BSAC and all other isolates was performed previously using Velvet and assembly statistics can be found in Turner *et al*., 2019. We also repeated the assembly using SPAdes (Bankevich *et al*., 2012) to confirm the regions surrounding prophage integration. We chose to focus on ST39 isolates (or closely related STs) and excluded those with highly diverse STs. As previously identified (Turner *et al*., 2019), these STs represent diverse genetic backgrounds and therefore do not reflect the population described here. Short read sequence data was mapped to the reference strain MGAS10750 using SMALT (https://www.sanger.ac.uk/science/tools/smalt) and core SNPs (excluding prophage regions) were extracted and used to generate a maximum likelihood phylogenetic tree using RAxML (Stamatakis 2014) with the GTR substitution model and 100 bootstraps. Regions of recombination were predicted using Gubbins analysis using the default parameters (Croucher *et al*., 2015). Annotated coding regions of each prophage within MGAS10750 were extracted and the DNA sequence of each was used to BLAST *de novo* assemblies. Gene presence was assigned with >98% match over 100% length.

### Prophage Induction

To determine prophage induction, overnight cultures of *S. pyogenes* were diluted 1:10 and grown to an OD 600nm of 0.3 before cultures were split into two 50mL aliquots. One aliquot was treated with 0.2 μg/mL mitomycin C and the second served as an untreated control. Cultures were grown for an additional 3 hours before the culture was pelleted by centrifugation and DNA was extracted. Bacterial DNA was extracted using the method of Pospiech and Neumann (Pospiech and Neumann, 1995). Isolates were also cultured exponentially from 1:10 dilutions of overnight cultures, and DNA extracted from aliquots at given time points. To detect excision and integration of each of the three prophages and SpyCIM4, specific sets of four primer combinations were designed (Supplementary Table 2). PCR was performed using each primer combination.

### RNA extraction and transcription detection

RNA was extracted from one M4_complete_ isolate and one M4_degraded_ isolate cultured exponentially for 3 hours using a using a hot acidic phenol method, as described previously (Reglinski *et al*., 2019). RNA samples were then DNase treated with Turbo DNase *free* (Ambion) for 30 minutes at 37°C and 5 µg was converted to cDNA by reverse transcription using Transcriptor reverse transcriptase (Roche) and random hexaoligos (Sigma-Aldrich). A corresponding RT-negative reaction was also performed for each sample whereby the reverse transcriptase was excluded as a control for contaminating genomic DNA. Transcription of *mutL* and *mutS* was detected using standard PCR with 100ng of cDNA (or RT-negative equivalent) and primers designed to detect each transcript individually (Supplementary Table 2).

## Acknowledgements

The authors would like to thank the British Society for Antimicrobial Chemotherapy for providing the *emm*4 isolates.

## Supplementary Data

**Supplementary Table 1**. Details of isolates used in this study.

Supplementary_Table_1.xlsx

**Supplementary Table 2.**
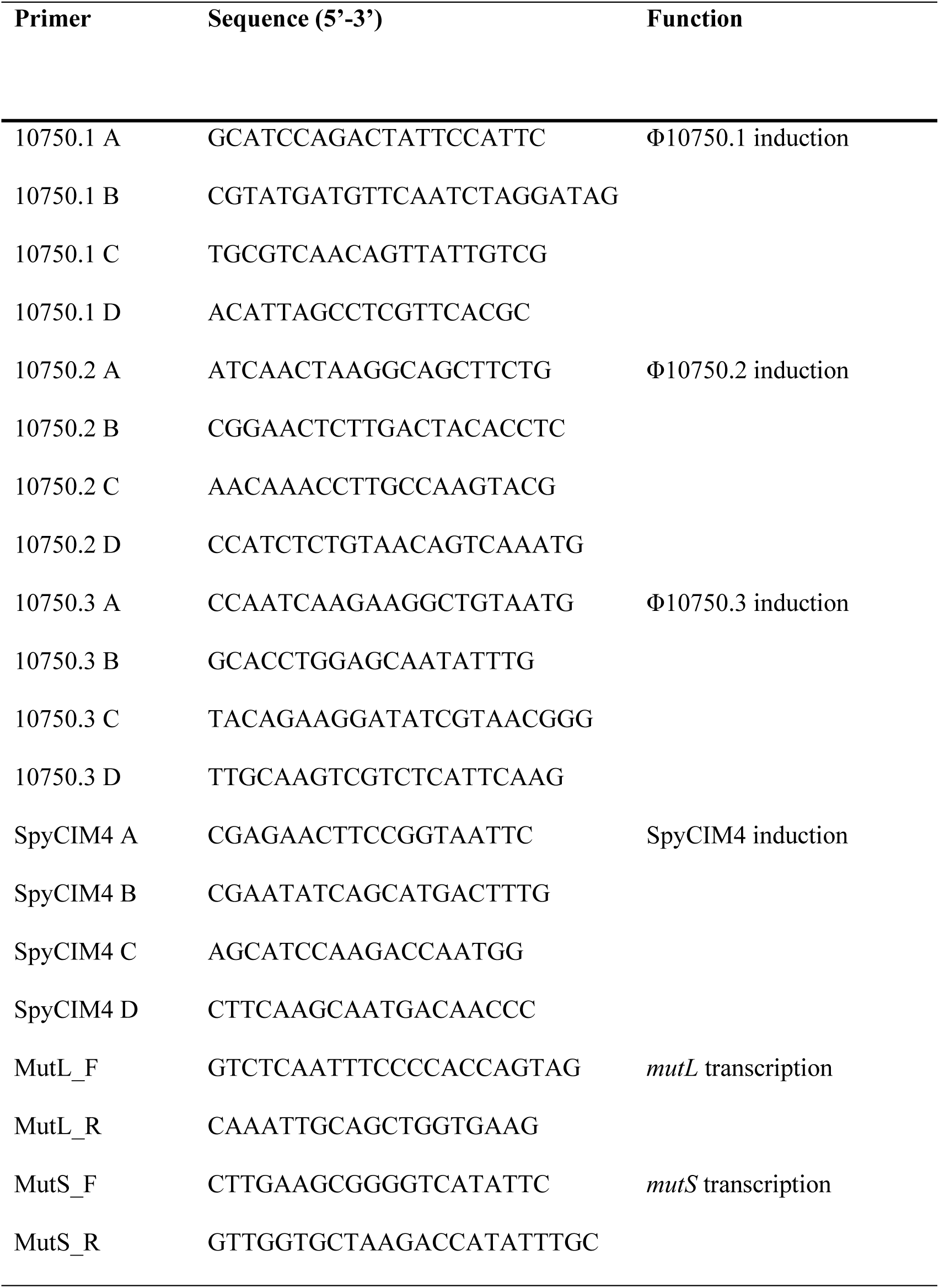
Primers used in the study.

## Supplementary Figures

**Supplementary Figure 1:**
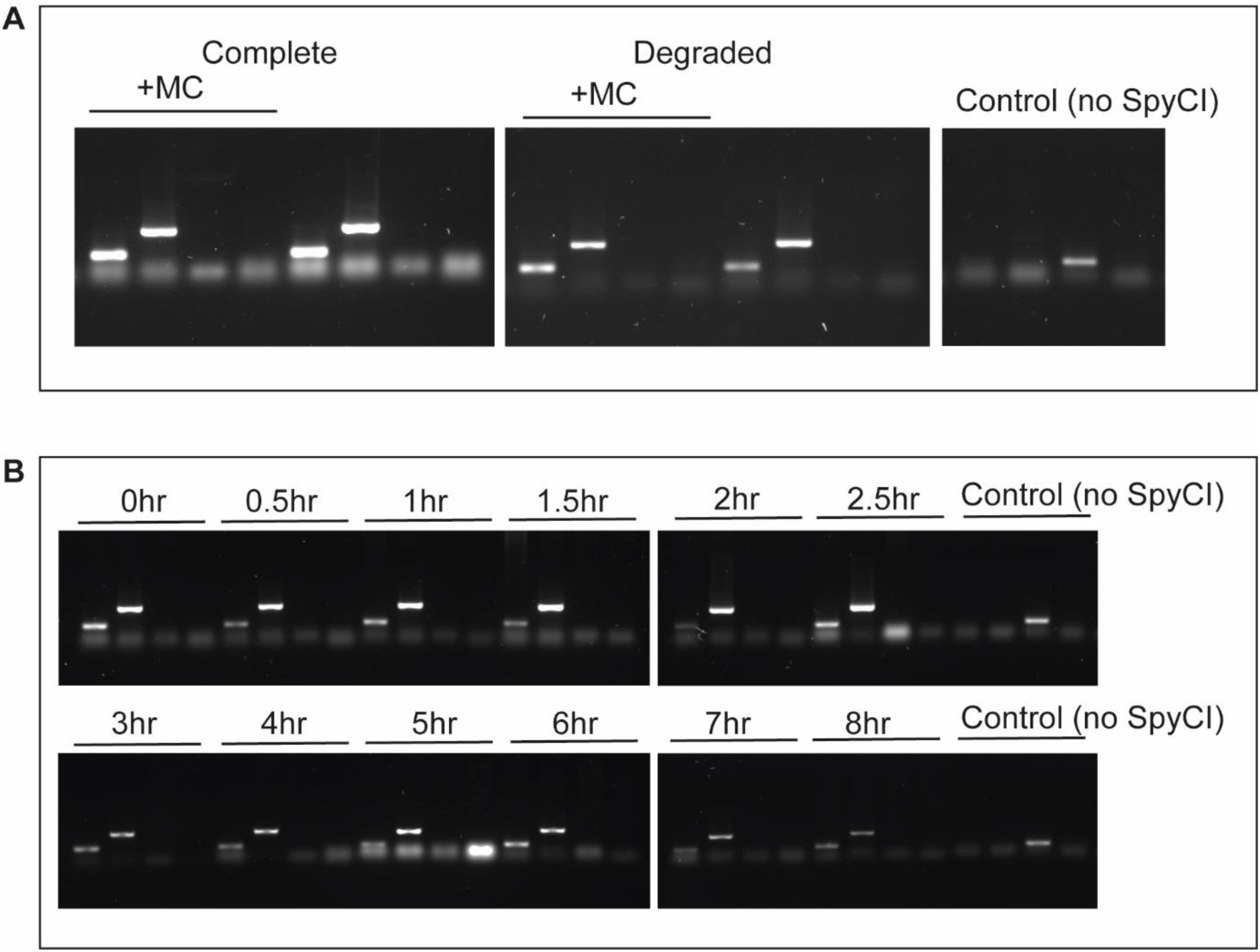
SpyCIM4 is not induced in any of the *emm*4 BSAC isolates. **A**. Primers were designed to detect integrated and excised SpyCI in genotype *emm*4 isolates in response to induction with mitomycin C. Only integrated SpyCI was detected (bands in first two lanes per sample) in both isolates, and no evidence of excised SpyCI (third and fourth lanes per sample), even in the presence of mitomycin C (+MC) **B**. To determine if the *emm*4 SpyCI would excise at specific stages in the bacterial growth curve, samples were taken at 30-minute intervals from 0-3 hours, then at 60-minute intervals thereafter, for a further 5 hours. Excised SpyCI was not detected at any point in bacterial growth in liquid culture. A control DNA sample was also used in each experiment extracted from *emm*89 strain H293; this strain has no SpyCI element and therefore only a band in third lane can be detected.

**Supplementary Figure 2.**
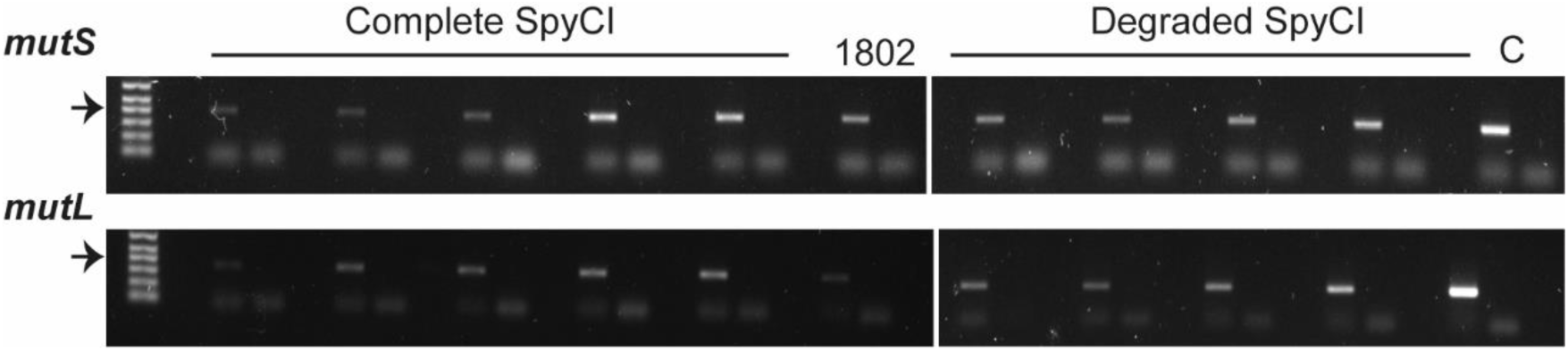
Expression of both *mutL* and *mutS* detected in both M4_complete_- and M4_degraded_ type *emm*4 isolates. Bacterial RNA was extracted from M4_complete_- and M4_degraded_ isolates at 3 hours of growth (early log) and converted to cDNA by reverse transcription (RT). 100ng of cDNA was used to detect *mutL* or *mutS* transcript by PCR. Each isolate is represented by two lanes; the first is the RT positive sample (ie containing cDNA), the second is a negative control whereby RT was excluded. Transcription of *mutS* (top gel) and *mutL* (bottom gel) was detected in all five BSAC M4_complete_ as well as all four BSAC M4_degraded_ isolates. Transcripts for *mutS* and *mutL* were also detected in BSAC_bs1802 which carries a different SpyCI element to the other *emm*4 strains. C; DNA from control SpyCI-negative H293 in the first lane, no-DNA sample in the second lane. Arrows indicate 400bp on the 1kb Plus Ladder (Invitrogen).

**Supplementary Figure 3.**
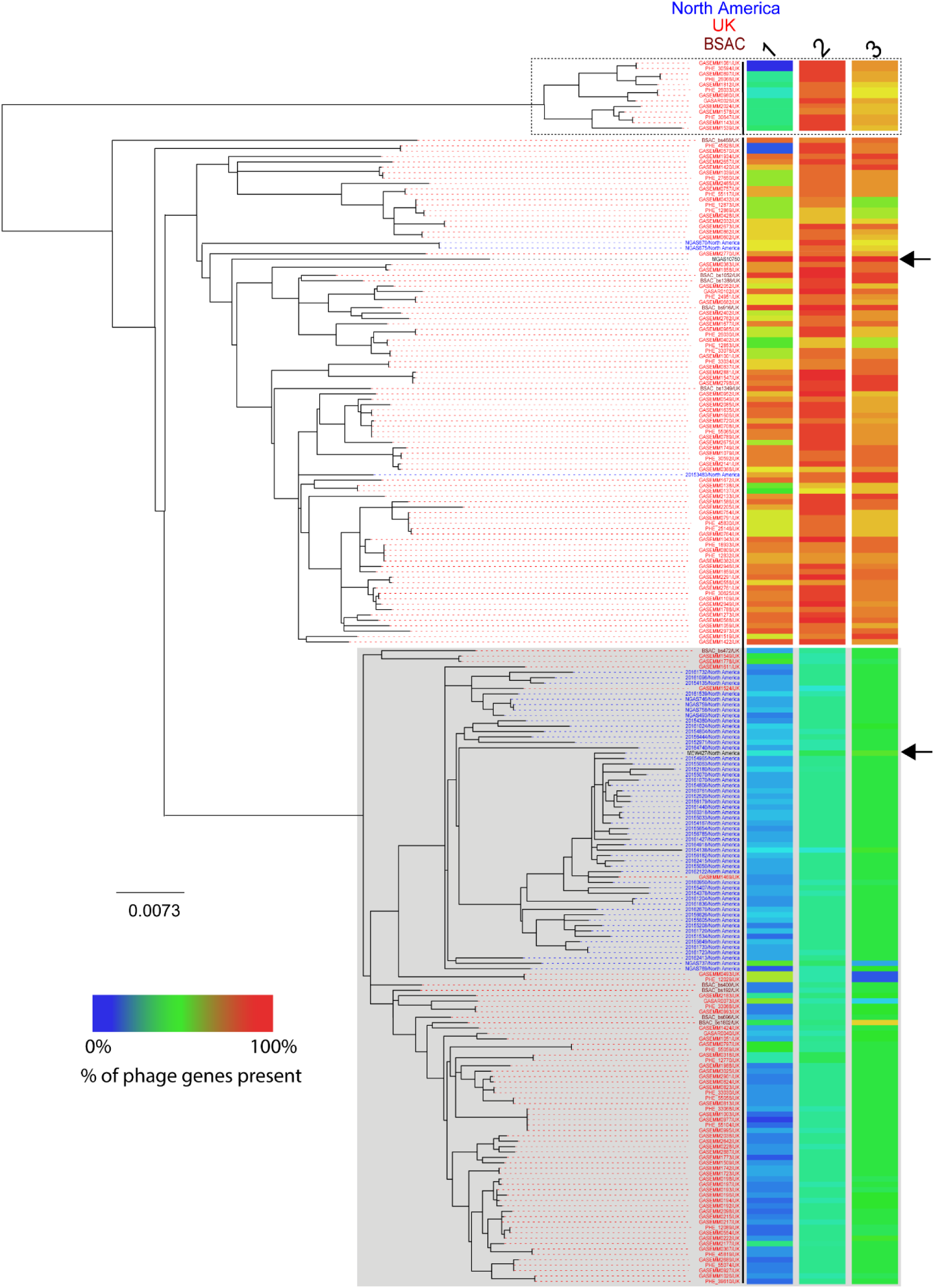
Lineages broadly associated with levels of prophage gene presence. Mid-rooted phylogenetic tree constructed from core SNPs obtained after mapping short read sequence data from 223 isolates as well as the reference genome MEW427 to MGAS10750 (as Figure 5). Lineages are defined as in Figure 5; MGAS10750-like (no shading), MEW427-like (grey shaded), separate 13 isolate lineage that differs by the first phage (dotted-line). The percentage (%) of genes present in the *de novo* assembled genome for each isolate was calculated by BLAST analysis for each of the three MGAS10750 prophages (1-3) and indicated by colour scale (key indicated). Two isolates within the MGAS10750 lineage do not have the first prophage as indicated by the blue shading in the first column. Top arrow indicates MGAS10750 and the lower arrow indicates MEW427. Isolates names are given for each branch and colour coded based on collection; Blue; North American isolates from the USA n=48 isolated 2015 (Chochua *et al*., 2017), and Canada (n=8 isolated 2013 or unknown date Athey *et al*. 2014, Athey *et al*. 2016),. Red; UK isolates (PHE n=153 (Chalker *et al*. 2017; Kapatai *et al*., 2017) isolated 2014/2015, Cambridgeshire n=4, 2008-2012, (Turner *et al*. 2017)), Brown; BSAC isolates (n=10, 2001-2008). Scale bar represents substitutions per site.

## References

1. Afshar, B., Turner, C. E., Lamagni, T. L., Smith, K. C., Al-Shahib, A., Underwood, A., Holden, M., Efstratiou, A., & Sriskandan, S. (2017). Enhanced nasopharyngeal infection and shedding associated with an epidemic lineage of *emm*3 group A *Streptococcus*. Virulence, 8(7), 1390–1400. https://doi.org/10.1080/21505594.2017.1325070

2. Al-Shahib, A., Underwood, A., Afshar, B., Turner, C. E., Lamagni, T., Sriskandan, S., & Efstratiou, A. (2016). Emergence of a novel lineage containing a prophage in *emm*/M3 group A *Streptococcus* associated with upsurge in invasive disease in the UK. Microbial genomics, 2(6), e000059. https://doi.org/10.1099/mgen.0.000059

3. Athey, T. B., Teatero, S., Li, A., Marchand-Austin, A., Beall, B. W., & Fittipaldi, N. (2014). Deriving group A *Streptococcus* typing information from short-read whole-genome sequencing data. Journal of clinical microbiology, 52(6), 1871–1876. https://doi.org/10.1128/JCM.00029-14

4. Athey, T. B., Teatero, S., Sieswerda, L. E., Gubbay, J. B., Marchand-Austin, A., Li, A., Wasserscheid, J., Dewar, K., McGeer, A., Williams, D., & Fittipaldi, N. (2016). High Incidence of Invasive Group A *Streptococcus* Disease Caused by Strains of Uncommon *emm* Types in Thunder Bay, Ontario, Canada. Journal of clinical microbiology, 54(1), 83–92. https://doi.org/10.1128/JCM.02201-15

5. Balasubramanian, S., Osburne, M. S., BrinJones, H., Tai, A. K., & Leong, J. M. (2019). Prophage induction, but not production of phage particles, is required for lethal disease in a microbiome-replete murine model of enterohemorrhagic *E. coli* infection. PLoS pathogens, 15(1), e1007494. https://doi.org/10.1371/journal.ppat.1007494

6. Bankevich A, Nurk S, Antipov D, Gurevich AA, Dvorkin M, Kulikov AS, Lesin VM, Nikolenko SI, Pham S, Prjibelski AD, Pyshkin AV, Sirotkin AV, Vyahhi N, Tesler G, Alekseyev MA, Pevzner PA. SPAdes: A New Genome Assembly Algorithm and Its Applications to Single-Cell Sequencing (2012). Journal of Computational Biology, 19, 455–477. doi: 10.1089/cmb.2012.0021

7. Banks, D. J., Beres, S. B., & Musser, J. M. (2002). The fundamental contribution of phages to GAS evolution, genome diversification and strain emergence. Trends in microbiology, 10(11), 515–521. https://doi.org/10.1016/s0966-842x(02)02461-7

8. Banks, D. J., Lei, B., & Musser, J. M. (2003). Prophage induction and expression of prophage-encoded virulence factors in group A *Streptococcus* serotype M3 strain MGAS315. Infection and immunity, 71(12), 7079–7086. https://doi.org/10.1128/iai.71.12.7079-7086.2003

9. Ben Zakour, N. L., Davies, M. R., You, Y., Chen, J. H., Forde, B. M., Stanton-Cook, M., Yang, R., Cui, Y., Barnett, T. C., Venturini, C., Ong, C. L., Tse, H., Dougan, G., Zhang, J., Yuen, K. Y., Beatson, S. A., & Walker, M. J. (2015). Transfer of scarlet fever-associated elements into the group A *Streptococcus* M1T1 clone. Scientific reports, 5, 15877. https://doi.org/10.1038/srep15877

10. Beres, S. B., & Musser, J. M. (2007). Contribution of exogenous genetic elements to the group A *Streptococcus* metagenome. PloS one, 2(8), e800. https://doi.org/10.1371/journal.pone.0000800

11. Beres, S. B., Richter, E. W., Nagiec, M. J., Sumby, P., Porcella, S. F., DeLeo, F. R., & Musser, J. M. (2006). Molecular genetic anatomy of inter- and intraserotype variation in the human bacterial pathogen group A *Streptococcus*. Proceedings of the National Academy of Sciences of the United States of America, 103(18), 7059–7064. https://doi.org/10.1073/pnas.0510279103

12. Beres, S. B., Sylva, G. L., Sturdevant, D. E., Granville, C. N., Liu, M., Ricklefs, S. M., Whitney, A. R., Parkins, L. D., Hoe, N. P., Adams, G. J., Low, D. E., DeLeo, F. R., McGeer, A., & Musser, J. M. (2004). Genome-wide molecular dissection of serotype M3 group A *Streptococcus* strains causing two epidemics of invasive infections. Proceedings of the National Academy of Sciences of the United States of America, 101(32), 11833–11838. https://doi.org/10.1073/pnas.0404163101

13. Broudy, T. B., Pancholi, V., & Fischetti, V. A. (2001). Induction of lysogenic bacteriophage and phage-associated toxin from group A streptococci during coculture with human pharyngeal cells. Infection and immunity, 69(3), 1440–1443. https://doi.org/10.1128/IAI.69.3.1440-1443.2001

14. Broudy, T. B., Pancholi, V., & Fischetti, V. A. (2002). The *in vitro* interaction of *Streptococcus pyogenes* with human pharyngeal cells induces a phage-encoded extracellular DNase. Infection and immunity, 70(6), 2805–2811. https://doi.org/10.1128/iai.70.6.2805-2811.2002

15. Canchaya, C., Proux, C., Fournous, G., Bruttin, A., & Brüssow, H. (2003). Prophage genomics. Microbiology and molecular biology reviews: MMBR, 67(2), 238–276. https://doi.org/10.1128/mmbr.67.2.238-276.2003

16. Carapetis, J. R., Steer, A. C., Mulholland, E. K., & Weber, M. (2005). The global burden of group A streptococcal diseases. The Lancet. Infectious diseases, 5(11), 685–694. https://doi.org/10.1016/S1473-3099(05)70267-X

17. Chalker, V., Jironkin, A., Coelho, J., Al-Shahib, A., Platt, S., Kapatai, G., Daniel, R., Dhami, C., Laranjeira, M., Chambers, T., Guy, R., Lamagni, T., Harrison, T., Chand, M., Johnson, A. P., Underwood, A., & Scarlet Fever Incident Management Team (2017). Genome analysis following a national increase in Scarlet Fever in England 2014. BMC genomics, 18(1), 224. https://doi.org/10.1186/s12864-017-3603-z

18. Chochua, S., Metcalf, B. J., Li, Z., Rivers, J., Mathis, S., Jackson, D., Gertz, R. E., Jr, Srinivasan, V., Lynfield, R., Van Beneden, C., McGee, L., & Beall, B. (2017). Population and Whole Genome Sequence Based Characterization of Invasive Group A Streptococci Recovered in the United States during 2015. mBio, 8(5), e01422–17. https://doi.org/10.1128/mBio.01422-17

19. Commons, R. J., Smeesters, P. R., Proft, T., Fraser, J. D., Robins-Browne, R., & Curtis, N. (2014). Streptococcal superantigens: categorization and clinical associations. Trends in molecular medicine, 20(1), 48–62. https://doi.org/10.1016/j.molmed.2013.10.004

20. Croucher, N. J., Page, A. J., Connor, T. R., Delaney, A. J., Keane, J. A., Bentley, S. D., Parkhill, J., & Harris, S. R. (2015). Rapid phylogenetic analysis of large samples of recombinant bacterial whole genome sequences using Gubbins. Nucleic acids research, 43(3), e15. https://doi.org/10.1093/nar/gku1196

21. Davies, M. R., Holden, M. T., Coupland, P., Chen, J. H., Venturini, C., Barnett, T. C., Zakour, N. L., Tse, H., Dougan, G., Yuen, K. Y., & Walker, M. J. (2015). Emergence of scarlet fever *Streptococcus pyogenes emm*12 clones in Hong Kong is associated with toxin acquisition and multidrug resistance. Nature genetics, 47(1), 84–87. https://doi.org/10.1038/ng.3147

22. DebRoy, S., Li, X., Kalia, A., Galloway-Pena, J., Shah, B. J., Fowler, V. G., Flores, A. R., & Shelburne, S. A. (2018). Identification of a chimeric *emm* gene and novel *emm* pattern in currently circulating strains of *emm*4 Group A *Streptococcus*. Microbial genomics, 4(11), e000235. https://doi.org/10.1099/mgen.0.000235

23. Desiere, F., Mcshan, W., Sinderen, D. V., Ferretti, J. J., & Brüssow, H. (2001). Comparative Genomics Reveals Close Genetic Relationships between Phages from Dairy Bacteria and Pathogenic Streptococci: Evolutionary Implications for Prophage-Host Interactions. Virology, 288(2), 325–341. doi: 10.1006/viro.2001.1085

24. Efstratiou, A., & Lamagni, T. (2017). Epidemiology of *Streptococcus pyogenes*. In Streptococcus pyogenes: basic biology to clinical manifestations [Internet]. University of Oklahoma Health Sciences Center.

25. Facklam, R., Beall, B., Efstratiou, A., Fischetti, V., Johnson, D., Kaplan, E., Kriz, P., Lovgren, M., Martin, D., Schwartz, B., Totolian, A., Bessen, D., Hollingshead, S., Rubin, F., Scott, J., & Tyrrell, G. (1999). *emm* typing and validation of provisional M types for group A streptococci. Emerging infectious diseases, 5(2), 247–253. https://doi.org/10.3201/eid0502.990209

26. Flores, A. R., Chase McNeil, J., Shah, B., Van Beneden, C., & Shelburne, S. A. (2019). Capsule-Negative *emm* Types Are an Increasing Cause of Pediatric Group A Streptococcal Infections at a Large Pediatric Hospital in Texas. Journal of the Pediatric Infectious Diseases Society, 8(3), 244–250. https://doi.org/10.1093/jpids/piy053

27. Flores, A. R., Jewell, B. E., Fittipaldi, N., Beres, S. B., & Musser, J. M. (2012). Human disease isolates of serotype m4 and m22 group A *Streptococcus* lack genes required for hyaluronic acid capsule biosynthesis. mBio, 3(6), e00413–12. https://doi.org/10.1128/mBio.00413-12

28. Frost, H. R., Davies, M. R., Delforge, V., Lakhloufi, D., Sanderson-Smith, M., Srinivasan, V., Steer, A. C., Walker, M. J., Beall, B., Botteaux, A., & Smeesters, P. R. (2020). Analysis of Global Collection of Group A *Streptococcus* Genomes Reveals that the Majority Encode a Trio of M and M-Like Proteins. mSphere, 5(1), e00806–19. https://doi.org/10.1128/mSphere.00806-19

29. Galloway-Peña, J., Clement, M. E., Sharma Kuinkel, B. K., Ruffin, F., Flores, A. R., Levinson, H., Shelburne, S. A., Moore, Z., & Fowler, V. G., Jr (2016). Application of Whole-Genome Sequencing to an Unusual Outbreak of Invasive Group A Streptococcal Disease. Open forum infectious diseases, 3(1), ofw042. https://doi.org/10.1093/ofid/ofw042

30. Hendrickson, C., Euler, C. W., Nguyen, S. V., Rahman, M., McCullor, K. A., King, C. J., Fischetti, V. A., & McShan, W. M. (2015). Elimination of Chromosomal Island SpyCIM1 from *Streptococcus pyogenes* Strain SF370 Reverses the Mutator Phenotype and Alters Global Transcription. PloS one, 10(12), e0145884. https://doi.org/10.1371/journal.pone.0145884

31. Ingrey, K. T., Ren, J., & Prescott, J. F. (2003). A fluoroquinolone induces a novel mitogen-encoding bacteriophage in *Streptococcus canis*. Infection and immunity, 71(6), 3028–3033. https://doi.org/10.1128/iai.71.6.3028-3033.2003

32. Jacob, K. M., Spilker, T., LiPuma, J. J., Dawid, S. R., & Watson, M. E., Jr (2016). Complete Genome Sequence of *emm*4 *Streptococcus pyogenes* MEW427, a Throat Isolate from a Child Meeting Clinical Criteria for Pediatric Autoimmune Neuropsychiatric Disorders Associated with Streptococcus (PANDAS). Genome announcements, 4(2), e00127–16. https://doi.org/10.1128/genomeA.00127-16

33. Jaggi, P., Tanz, R. R., Beall, B., & Shulman, S. T. (2005). Age influences the *emm* type distribution of pediatric group A streptococcal pharyngeal isolates. The Pediatric infectious disease journal, 24(12), 1089–1092. https://doi.org/10.1097/01.inf.0000190023.89759.96

34. Kapatai, G., Coelho, J., Platt, S., & Chalker, V. J. (2017). Whole genome sequencing of group A *Streptococcus*: development and evaluation of an automated pipeline for *emm* gene typing. PeerJ, 5, e3226. https://doi.org/10.7717/peerj.3226

35. Kim, S., Lee, S., Park, H., & Kim, S. (2019). Predominance of *emm*4 and antibiotic resistance of *Streptococcus pyogenes* in acute pharyngitis in a southern region of Korea. Journal of medical microbiology, 68(7), 1053–1058. https://doi.org/10.1099/jmm.0.001005

36. Luca-Harari, B., Darenberg, J., Neal, S., Siljander, T., Strakova, L., Tanna, A., Creti, R., Ekelund, K., Koliou, M., Tassios, P. T., van der Linden, M., Straut, M., Vuopio-Varkila, J., Bouvet, A., Efstratiou, A., Schalén, C., Henriques-Normark, B., Strep-EURO Study Group, & Jasir, A. (2009). Clinical and microbiological characteristics of severe *Streptococcus pyogenes* disease in Europe. Journal of clinical microbiology, 47(4), 1155–1165. https://doi.org/10.1128/JCM.02155-08

37. Maiques, E., Ubeda, C., Campoy, S., Salvador, N., Lasa, I., Novick, R. P., Barbé, J., & Penadés, J. R. (2006). Beta-lactam antibiotics induce the SOS response and horizontal transfer of virulence factors in *Staphylococcus aureus*. Journal of bacteriology, 188(7), 2726–2729. https://doi.org/10.1128/JB.188.7.2726-2729.2006

38. McMillan, D. J., Drèze, P. A., Vu, T., Bessen, D. E., Guglielmini, J., Steer, A. C., Carapetis, J. R., Van Melderen, L., Sriprakash, K. S., & Smeesters, P. R. (2013). Updated model of group A *Streptococcus* M proteins based on a comprehensive worldwide study. Clinical microbiology and infection: the official publication of the European Society of Clinical Microbiology and Infectious Diseases, 19(5), E222–E229. https://doi.org/10.1111/1469-0691.12134

39. McShan, W. M., McCullor, K. A., & Nguyen, S. V. (2019). The Bacteriophages of *Streptococcus pyogenes*. Microbiology spectrum, 7(3), 10.1128/microbiolspec.GPP3-0059-2018. https://doi.org/10.1128/microbiolspec.GPP3-0059-2018

40. Nakagawa, I., Kurokawa, K., Yamashita, A., Nakata, M., Tomiyasu, Y., Okahashi, N., Kawabata, S., Yamazaki, K., Shiba, T., Yasunaga, T., Hayashi, H., Hattori, M., & Hamada, S. (2003). Genome sequence of an M3 strain of *Streptococcus pyogenes* reveals a large-scale genomic rearrangement in invasive strains and new insights into phage evolution. Genome research, 13 (6A), 1042–1055. https://doi.org/10.1101/gr.1096703

41. Nasser, W., Beres, S. B., Olsen, R. J., Dean, M. A., Rice, K. A., Long, S. W., Kristinsson, K. G., Gottfredsson, M., Vuopio, J., Raisanen, K., Caugant, D. A., Steinbakk, M., Low, D. E., McGeer, A., Darenberg, J., Henriques-Normark, B., Van Beneden, C. A., Hoffmann, S., & Musser, J. M. (2014). Evolutionary pathway to increased virulence and epidemic group A *Streptococcus* disease derived from 3,615 genome sequences. Proceedings of the National Academy of Sciences of the United States of America, 111(17), E1768–E1776. https://doi.org/10.1073/pnas.1403138111

42. Pospiech, A. (1995). A versatile quick-prep of genomic DNA from Gram-positive bacteria. Trends in Genetics, 11 (6), 217–218. doi: https://doi.org/10.1016/S0168-9525(00)89052-6

43. Ramisetty, B., & Sudhakari, P. A. (2019). Bacterial ‘Grounded’ Prophages: Hotspots for Genetic Renovation and Innovation. Frontiers in genetics, 10, 65. https://doi.org/10.3389/fgene.2019.00065

44. Reglinski, M., Sriskandan, S., & Turner, C. E. (2019). Identification of two new core chromosome-encoded superantigens in *Streptococcus pyogenes; speQ* and *speR*. Journal of Infection, 78(5), 358–363. doi: 10.1016/j.jinf.2019.02.005

45. Remmington A, Turner CE. The DNases of pathogenic Lancefield streptococci. Microbiology. 2018;164(3):242–250. doi:10.1099/mic.0.000612

46. Scott, J., Nguyen, S. V., King, C. J., Hendrickson, C., & McShan, W. M. (2012). Phage-Like *Streptococcus pyogenes* Chromosomal Islands (SpyCI) and Mutator Phenotypes: Control by Growth State and Rescue by a SpyCI-Encoded Promoter. Frontiers in microbiology, 3, 317. https://doi.org/10.3389/fmicb.2012.00317

47. Scott, J., Thompson-Mayberry, P., Lahmamsi, S., King, C. J., & McShan, W. M. (2008). Phage-associated mutator phenotype in group A *Streptococcus*. Journal of bacteriology, 190(19), 6290–6301. https://doi.org/10.1128/JB.01569-07

48. Silva-Costa, C., Carriço, J. A., Ramirez, M., & Melo-Cristino, J. (2014). Scarlet fever is caused by a limited number of *Streptococcus pyogenes* lineages and is associated with the exotoxin genes *ssa, speA* and *speC*. The Pediatric infectious disease journal, 33(3), 306–310. https://doi.org/10.1097/INF.0000000000000088

49. Stamatakis A. (2014). RAxML version 8: a tool for phylogenetic analysis and post-analysis of large phylogenies. Bioinformatics (Oxford, England), 30(9), 1312–1313. https://doi.org/10.1093/bioinformatics/btu033

50. Tse, H., Bao, J. Y., Davies, M. R., Maamary, P., Tsoi, H. W., Tong, A. H., Ho, T. C., Lin, C. H., Gillen, C. M., Barnett, T. C., Chen, J. H., Lee, M., Yam, W. C., Wong, C. K., Ong, C. L., Chan, Y. W., Wu, C. W., Ng, T., Lim, W. W., Tsang, T. H., … Yuen, K. Y. (2012). Molecular characterization of the 2011 Hong Kong scarlet fever outbreak. The Journal of infectious diseases, 206(3), 341–351. https://doi.org/10.1093/infdis/jis362

51. Turner, C. E., Abbott, J., Lamagni, T., Holden, M. T., David, S., Jones, M. D., Game, L., Efstratiou, A., & Sriskandan, S. (2015). Emergence of a New Highly Successful Acapsular Group A *Streptococcus* Clade of Genotype *emm*89 in the United Kingdom. mBio, 6(4), e00622. https://doi.org/10.1128/mBio.00622-15

52. Turner, C. E., Pyzio, M., Song, B., Lamagni, T., Meltzer, M., Chow, J. Y., Efstratiou, A., Curtis, S., & Sriskandan, S. (2016). Scarlet Fever Upsurge in England and Molecular-Genetic Analysis in North-West London, 2014. Emerging infectious diseases, 22(6), 1075–1078. https://doi.org/10.3201/eid2206.151726

53. Turner, C. E., Holden, M., Blane, B., Horner, C., Peacock, S. J., & Sriskandan, S. (2019). The Emergence of Successful *Streptococcus pyogenes* Lineages through Convergent Pathways of Capsule Loss and Recombination Directing High Toxin Expression. mBio, 10(6), e02521–19. https://doi.org/10.1128/mBio.02521-19

54. Turner, C. E., Bedford, L., Brown, N. M., Judge, K., Török, M. E., Parkhill, J., & Peacock, S. J. (2017). Community outbreaks of group A *Streptococcus* revealed by genome sequencing. Scientific reports, 7(1), 8554. https://doi.org/10.1038/s41598-017-08914-x

55. Whitehead, B. D., Smith, H. V., & Nourse, C. (2011). Invasive group A streptococcal disease in children in Queensland. Epidemiology and infection, 139(4), 623–628. https://doi.org/10.1017/S0950268810001378

56. Yan, J. J., Liu, C. C., Ko, W. C., Hsu, S. Y., Wu, H. M., Lin, Y. S., Lin, M. T., Chuang, W. J., & Wu, J. J. (2003). Molecular analysis of group A streptococcal isolates associated with scarlet fever in southern Taiwan between 1993 and 2002. Journal of clinical microbiology, 41(10), 4858–4861. https://doi.org/10.1128/jcm.41.10.4858-4861.2003

57. Zhang, X., McDaniel, A. D., Wolf, L. E., Keusch, G. T., Waldor, M. K., & Acheson, D. W. (2000). Quinolone antibiotics induce Shiga toxin-encoding bacteriophages, toxin production, and death in mice. The Journal of infectious diseases, 181(2), 664–670. https://doi.org/10.1086/315239

